# Invasive aliens threatened with native extinction: examining best practice for species translocations under climate change

**DOI:** 10.1101/429084

**Authors:** Paul A. Egan, David Bourke, Wilfried Thuiller, Maude E.A. Baudraz, Damien Georges, Julien Renaud, Jane C. Stout

**Affiliations:** Swedish University of Agricultural Sciences, Department of Plant Protection Biology, PO Box 102, SE-230 53 Alnarp, Sweden; Department of Botany, School of Natural Sciences, Trinity College Dublin, Dublin 2, Ireland; School of Natural Sciences, Liverpool John Moores University, Liverpool L3 3AF, UK; Univ. Grenoble Alpes, CNRS, Univ. Savoie Mont-Blanc, Laboratoire d’Ecologie Alpine (LECA), F-38000 Grenoble, France; Department of Zoology, School of Natural Sciences, Trinity College Dublin, Dublin 2, Ireland

**Keywords:** assisted colonization, assisted migration, climate change, invasive alien species, microrefugia, narrow endemic species, species distribution modelling, translocation

## Abstract

**Aim:** Translocation remains a controversial strategy in species conservation. Here, we utilise the unusual scenario of invasive alien species (IAS) threatened with extinction in their native range to address key challenges in deciding ‘whether’, ‘where’, and ‘when’ to implement translocation, and how best to approach conservation under seemingly contradictory circumstances.

**Location:** Iberian Peninsula, NW Europe

**Methods:** *Rhododendron ponticum* ssp. *baeticum* was selected as a model IAS for case study analysis. We used species distribution models (SDMs) coupled with dynamic simulations of migration to assess: 1. the extinction risk posed to this species in its native Iberian range under climate change; 2. whether SDMs calibrated on the native range (replicating typical translocation planning) could predict invasive capacity in NW Europe; and 3. the extent to which recommended biogeographical constrains on translocations may limit available options. Insights gained on the above were used to build and test a generic decision framework for translocation, based on robust identification of microrefugia.

**Results:** Our findings suggest a high likelihood of climate-induced extinction for *R.p*. ssp. *baeticum* in its native range. Notably, SDMs completely failed to predict invasive capacity in NW Europe. However, application of our framework was successful in identifying sites more proximate to the native range – albeit outside this species’ current biogeographic region – potentially suited to translocation.

**Main conclusions:** The framework here developed can be used to guide translocation of climate-endangered species in a spatially and temporally precise manner. However, we caution that use of SDMs can possess short-comings in failing to capture a full picture of sites suited to translation, and in risk assessment of the capacity of translocated taxa to form invasive species. Strict biogeographic constraints to the selection of translocation sites can evidently help to safeguard against invasions, but may also severely hinder the options available to avert climate-induced extinctions.

## INTRODUCTION

Translocation, the intentional introduction of threatened species into a given geographic region, may be one of the few viable solutions available to avert species extinctions under rapid climate change (Hunter, 2007; Ledig et al., 2012; Gallagher et al., 2015). However, considerable controversy surrounds the selection of geographic areas for the purpose of translocation (Ricciardi & Simberloff, 2009b; IUCN/SSC, 2013). For instance, when species are translocated into an area within their historical distribution, this is referred to as a ‘reintroduction’, but if moved to a novel location, this is referred to as ‘assisted colonization’ (Seddon, 2010; Hällfors et al., 2014). Straightforward cases involve reintroduction of a species into areas of recent former occupancy where local extinction is directly attributed to anthropogenic influences (e.g. Baker et al., 2011; Nussear et al., 2012), or the assisted colonisation of a species into suitable neighbouring habitats (Seddon et al., 2015). In more complex cases, however, translocation outside of a species’ historical range may offer the only realistic prospect for curtailing extinctions (Thomas, 2011).

Although translocation outside of a species’ historical range has been justified on hypothetical grounds, few practical examples have emerged to date (Seddon et al., 2015), limiting insight into the technical processes and risks associated with this strategy (McLachlan et al., 2007; Hancock & Gallagher, 2014). In particular, significant challenges are presented in the form of risk assessment of translocation outside of a species’ historical range (such as the potential for aggressive invasion), and in the geographic identification of suitable sites (Mueller & Hellmann, 2008; Gallagher et al., 2015; Seddon et al., 2015). As such, and coupled with their high associated monetary costs, translocations (and in particular those within novel geographic areas) are generally considered the last line of action for averting species extinctions (Shoo et al., 2013; Kling et al., 2016).

In addition to the above challenges, identification of suitable sites for translocation prove further problematic in cases where: 1. a species’ historical range is unknown; 2. little to no suitable habitat remains within a known historical range (presently, or as projected under climate change); and 3. suitable translocation sites are not within close (bio)geographic proximity to the extant range. Such cases pose a challenge to proposed criteria for the appropriate use of translocation as a conservation strategy, which suggest that recipient site selection should be constrained to within a species’ continental and biogeographic region of occurrence (Hoegh-Guldberg et al., 2008; Thomas, 2011; Abeli et al., 2014). Restricting translocations to within such areas may therefore help to minimize invasive risk, as well as to preserve co-evolved biotic interactions. However, the potential ramifications that within-continent biogeographic constraints hold for translocation, in terms of limiting options to curtail extinctions, at present remain unclear and untested.

While the theoretical foundations which underpin translocation as a conservation strategy have been steadily refined over the last two decades (Seddon et al., 2015; Hällfors et al., 2017), further developments in the context of climate change remain an outstanding priority (Gallagher et al., 2015). In particular, recent elaboration of the concept of climate change refugia (Ashcroft, 2010; Keppel et al., 2012; Keppel et al., 2015; Lawler et al., 2015; Morelli et al., 2016) are of large potential relevance, and provide a theoretical basis for the identification of species-specific safe havens for biodiversity under future changing climate. Depending on the spatial and temporal dynamics of such safe havens, these may be termed as microrefugia, holdouts, or stepping-stones that facilitate species range shifts (Hannah et al., 2014). However, despite the potential to better guide translocation practise, explicit frameworks to support the selection of climate-stable habitat in the context of climate change remain in need of development (see for instance Alagador et al., 2014; Ferrarini et al., 2016; Payne & Bro-Jørgensen, 2016). Furthermore, although species distribution models (SDMs) are a standard recommended tool for the purpose of informing translocation (IUCN/SSC, 2013), and offer promising potential for the identification of climate change microrefugia (Ashcroft, 2010; McLaughlin & Zavaleta, 2012; Guisan et al., 2013), their use for such inference has to date been limited (Keppel et al., 2012), and seldom explicit to conservation translocation (Ferrarini et al., 2016; Hällfors et al., 2016; Hällfors et al., 2017).

As a means to address the key questions of ‘whether’, ‘where’, and ‘when’ to implement translocation, we collated examples of invasive alien species (IAS) which, paradoxically, are threatened with extinction in their native range due to climatic or anthropogenic factors (Suppl. Table 1). Given this juxtaposition, these species are useful to test the potential risks associated with translocation as a conservation strategy. In particular, we build a case-study on the Iberian tertiary relict plant *Rhododendron ponticum* ssp. *baeticum* (Boiss. & Reut.) Hand.-Mazz. (Ericaceae). Although highly invasive in NW Europe (Cross, 1975; Mejías et al., 2002; Mejías et al., 2007; Maclean et al., 2018), within its native range this species is subject to threats common to many climate-endangered taxa (e.g. Reinhardt et al., 2005; Early & Sax, 2011; Thomas, 2011), in that potential range shifts are constrained by high habitat specificity (within regionally rare or relict habitats), an inherent low dispersal ability, and the likely formation of insurmountable climatic barriers in the short to medium term.

Specifically, this study had three main aims. Firstly, we utilized spatially-explicit spread models (which coupled dynamic simulations of dispersal with projected shifts in climatic suitability from SDMs) to forecast the likelihood for successful migration and survival of native *R.p*. ssp. *baeticum* under climate change. Secondly, we examined to what extent biogeographic constraints may limit the options available for translocation of this species in Europe, and whether SDMs would have provided effective *a priori* risk assessment of this species’ invasive capacity in NW Europe. To replicate the information typically available for translocation planning and modelling, we hence restricted calibration of SDMs to within this species’ native Iberian range. Finally, based on this case-study, we developed and applied a generic framework to identify stable microrefugia potentially suited to translocation of climate-endangered species, accounting explicitly for climatic and statistical uncertainties. As a whole, insights gained from these investigations can contribute to last line efforts to curtail species extinctions under advancing climate change.

## METHODS

### Climate change data and scenarios

Baseline and future climate change datasets developed by the EU FP7-funded project EcoChange (Challenges in assessing and forecasting biodiversity and ecosystem changes in Europe) were used to undertake species distribution modelling at a 2 km resolution. This climate dataset was derived from Worldclim data (Hijmans et al., 2005) downscaled to a finer 100 m resolution. Further documentation of this dataset is available at crudata.uea.ac.uk/projects/ecochange/climatedata. For the current study, the EcoChange finer resolution raster data were aggregated to 2 km resolution using bilinear interpolation implemented in R. From an extensive set of environmental variables, six variables were selected for modelling (Suppl. Table 2), including three temperature variables, two precipitation variables, and potential evapotranspiration. This sub-set of variables was selected based on demonstrated performance in models, but also on their readily interpretable ecophysiological link to the fundamental niche of *R. ponticum* (Cross, 1975; Shaw, 1984; Griffin, 1994; Mejías et al., 2002; Mejías et al., 2007; Harris et al., 2009). Projections of future climatic change were based on the ECHAM5 general circulation model (GCM), which was coupled to the RCA30 regional climate model (Jones et al., 2004). Predictions were generated for two future time slices (2021–2050 and 2051–2080), and three different emissions scenarios (A1B, A2, and B2) based on the Special Report on Emissions Scenarios (SRES) from the IPCC’s Fourth Assessment Report.

These scenarios reflect different projections of demographic, socioeconomic, and technological development and their resultant influence on greenhouse gas emissions (van Vuuren et al., 2010). Although SRES scenarios have been more recently superseded by Representative Concentration Pathways (RCPs) in the IPCC’s Fifth Assessment Report, equivalences between these systems nonetheless exist. Hence up to 2080, the latest time period considered in this study, in increasing order of future projected CO_2_ concentrations; SRES scenario B2 closely resembles RCP4.5, A1B is virtually identical to RCP6.0, whereas A2 is intermediate between RCP6.0 and 8.5 (Moss et al., 2010; Meinshausen et al., 2011).

### Species distribution modelling

Spatial occurrence records for *R. ponticum* ssp. *baeticum* were assembled from field surveys conducted throughout 2011 and 2012 across the disjunct regions of this species’ native Iberian range (southern Spain, and southern and northern Portugal). Spatial coordinates were recorded to high (ca. 5 m) accuracy with a handheld GPS. Field records were combined with spatial coordinates obtained from literature sources (Erfmeier & Bruelheide, 2004; Perez Latorre & Cabezudo, 2006; Mejías et al., 2007). This combined dataset consisted of almost 200 unique occurrence points that were mostly non-aggregated in space (i.e. ≥ 50 m distance), and of resolution finer than the 2 km extent of climate data used here.

Current and future projected climatic suitability for *R. ponticum* ssp. *baeticum* was modelled using an ensemble of four regression and machine learning techniques (i.e. Random Forests (RFs), Generalised Boosting Models (GBMs), Generalised Additive Models (GAMs), and Maxent) available in the biomod2 library (Thuiller et al., 2009) and implemented within the R statistical programming environment (R Core Team, 2016). Default options were used for each model type (e.g. the number of trees used in RFs and GBMs). For the modelling techniques which required presence-absence data (all but Maxent, which required presence-only data), we used the random strategy in biomod2 to generate pseudo-absences at 10 times the number of presences in each model. Three evaluation runs were made for each model type, using three different sets of randomly generated pseudo-absences per run (affording a total of 36 replicates). Model evaluation was carried out by randomly splitting the data; 70% for calibration and 30% for validation, following which we assessed the area under the curve of the Receiver Operating Characteristic (ROC) (Swets, 1988) and the True Skill Statistic (TSS) (Allouche et al., 2006). All models were well supported (mean TSS = 0.989 ± 0.011 SD, mean ROC = 0.998 ± 0.003 SD, n = 36). All six environmental variables were included in each model run. Of these variables, precipitation of the driest quarter and mean temperature of the coldest quarter held the highest influence (Supp. Fig. 1), as assessed by the variable importance function in biomod2.

The specific limitations of SDMs are well noted (Guisan & Thuiller, 2005), and it is recognised that uncertainty associated with differences between SDM model techniques (i.e. statistical uncertainty), as well as future climate change scenarios (i.e. climatic uncertainty), can be considerable (Franklin et al., 2014). The development of an ‘ensemble’ approach to modelling, in which individual models are combined to produce a kind of meta model, provides one solution to account for statistical uncertainty (Araújo & New, 2007; Thuiller et al., 2013; Breiner et al., 2015). Here, we used a range of ensemble model (EM) techniques to combine the outputs from the four cross-validated SDMs; namely mean of probabilities (me); median of probabilities (med); committee averaging (ca); and weighted mean of probabilities (wm). Following evaluation of these four EMs calibrated on the native Iberian range (mean TSS = 0.992 ± 0.003 SD), these models were projected both in space (across Western Europe, including this species’ invasive range), and in time (across the current and future climate time slices considered). Final output included current and future predicted climatic suitability scores for each 2 km grid square, on both a continuous (0 to 1000) and binary (0 or 1) scale. Binary scores were converted from continuous scores based on a threshold value that optimised the TSS evaluation metric.

### Integrating dispersal constraints into SDMs

Predictions of future migration of *R.p*. ssp. *baeticum* under climate change were generated using the R package MigClim (Engler et al., 2012). Predictions were made using ensemble model spatial layers (as generated above), over which dynamic simulations of migration were implemented based on dispersal and life history parameters specific to this species (see below). Grid-based projections of dispersal were implemented at a cell resolution of 30 m. For this, SDM climatic suitability scores were therefore resampled through bilinear interpolation to match this grid resolution, and then converted into binary suitability/unsuitability values by specifying a classification threshold using ‘binary mode’ in MigClim. Data on dispersal and life history traits for parameterization of the MigClim model were obtained from available literature sources. For *R. ponticum*, these features included a negative exponential kernel with propagule dispersal up to 120 m (i.e. 4 grid cells) distance (Griffin, 1994; Harris et al., 2009; Travis et al., 2011); a minimum age of reproductive maturity at 11 years (Cross, 1975; Harris et al., 2009); followed by a logistic rate of increase in propagule production until fecundity levels-off around year 40 (Griffin, 1994). Occasional stochastic long-distance dispersal (LDD) events are suspected to occur for this species at distances which may exceed 1 km in turbulent air conditions (Shaw, 1984; Griffin, 1994), such as in mountainous regions.

LDD was therefore set in the range of 150–1500 m, at a frequency of 2% once full propagule production potential had been obtained. We purposely excluded the addition of landscape barriers to these models (i.e. any land cover type not potentially colonisable, such as agricultural land, urban areas, grassland etc.). This was done in order to simulate the most favourable circumstance in which all land cover types, if possessing suitable climate, could be colonized once dispersed into. Output was generated for the most optimistic combination of climate change scenario and ensemble model (B1 – committee averaging), as well for the most pessimistic scenario of climate change and ensemble model (A2 – mean averaging). For both scenarios, results are presented for each time period (2021–2050 and 2051–2080) and for each native disjunct region (southern Spain, northern and southern Portugal).

### A framework to identify stable microrefugia must be robust to uncertainties

As a test of a developed decision framework for translocation under climate change (Fig. 1.), we sought to identify stable microrefugia for *R.p*. ssp. *baeticum* based on a high level of accordance in suitability across climate change scenarios (B1, A1B, A2), time slices (2021–2050, 2051–2080), and ensemble models (mean, weighed mean, median, committee averaging). As input to the framework (Fig. 1.), cell climatic suitability scores were hence summed across these 24 SDM binary layers. Owing to these layer properties, identification of putative microrefugia was hence robust, in that statistical and climatic uncertainties were explicitly accounted for. Rasters of the summed individual cell suitability values (0 or 1) were generated, in which cells which possessed high accordance between layers thus exhibited high scores (to a maximum of 24). We here defined a cut-off of ≥ 20 (or 83.3 % accordance) as a reasonable threshold of robustness to uncertainties. Alternative approaches to the above work flow are also possible, depending on the type of score (binary vs. continuous) and model (ensemble vs. cross-validated models) used to identify cells possessing high accordance in predicted suitability. High scoring cells identified by the above process were further subject to an aggregation criterion, which restricted the selection of potential translocation sites to those which were spatially clustered into groups of 3 or more adjoining cells (i.e. ≥ 36 km^2^ total contiguous area). We deemed this area to be a minimum desired range size for translocation of *R.p*. ssp. *baeticum* based on ecological considerations. The final output of aggregated cells represented identified putative microrefugia, and were converted into polygon shapefiles.

**Figure 1.**
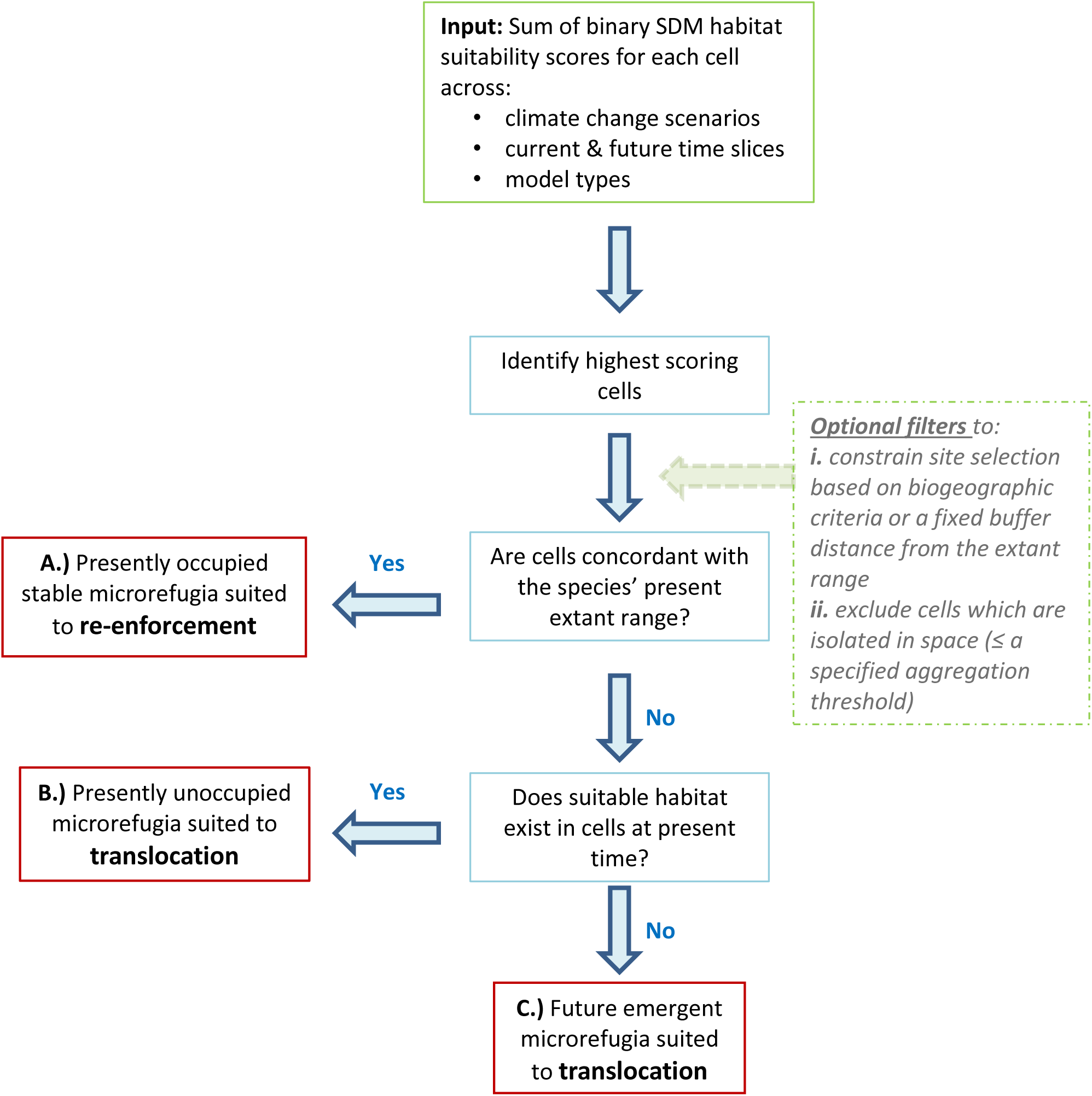
A decision framework for identification of robust, climate-stable microrefugia potentially suited to translocation, and associated management options. Identification of sites with high accordance in predicted suitability across the various climate change scenarios and model types can help guide decision-making by taking into account the risk of uncertainties. Given the temporal and spatially explicit nature of this framework, informed decisions can be made on ‘whether’, ‘where’ and ‘when’ translocation should be implemented.

Based on the concordance between putative microrefugia and the present extant occurrence of a species, decisions on ‘whether’, ‘where’ and ‘when’ to initiate translocation can potentially be made. Any partial overlap between identified microrefugia and a species’ extant range therefore signifies that at least some future stable sites are presently occupied (Fig. 1A); the most ideal scenario for conservation. However, if a given species is absent from identified microrefugia, such sites may represent strong candidates for immediate translocation (Fig. 1B). Whereas if a given species is absent from sites in which microrefugia will only emerge in the near future, such sites may represent good candidates for future translocation upon their eventual emergence (Fig. 1C). Based on the application of this framework, it is hence possible that none to all three microrefugia types (Fig. 1 A-C) can be simultaneously identified for a given species. As per Ferrarini *et al*. (2016), ex-situ conservation may be deemed the most reasonable course of action where no or little future suitable climate is identified across a geographic area of interest.

## RESULTS

### Dispersal-limited migration under climate change

Under the most optimistic scenario of climate change and ensemble model combination (B1 – committee averaging) employed, areas of suitable climate for *Rhododendron ponticum* ssp. *baeticum* was predicted to have significantly contracted by 2080 in one of its three disjunct ranges (southern Portugal), and have disappeared completely in another (southern Spain) (Table 1). While a limited amount of dispersal into areas of future suitable climate was predicted as possible in northern and southern Portugal; the vast majority of future suitable climate was predicted as non-accessible for dispersal and colonisation by the species (Table 1). In southern Spain, it was forecast that by 2050 the species would no longer occupy any area possessing suitable climate.

**Table 1.** Predicted climatic suitability and dispersal-limited migration of *R. ponticum* ssp. *baeticum* under climate change. Results indicate the progressive number of cells (at a spatial resolution of 30 m) which possess climatic suitability within this species’ three disjunct native Iberian regions. The simulation was conducted for both the most optimistic (B1 scenario/ ca EM) and pessimistic (A2 scenario/ mean EM) climate change scenario and ensemble model combination. Cells were enumerated as ‘total suitable’ (no. cells which possessed climatic suitability), ‘occupied under no dispersal’ (progressive no. cells in which the species is predicted to occur if no successful dispersal were possible), and ‘occupied under partial dispersal’ (progressive no. cells in which the species is predicted to occur under ecologically feasible dispersal).

| Range | Period | Optimistic scenario<br>(no. cells) |  |  | Pessimistic scenario<br>(no. cells) |  |  |
| --- | --- | --- | --- | --- | --- | --- | --- |
|  |  | Total<br>suitable | Occupied<br>– no<br>dispersal | Occupied<br>– partial<br>dispersal | Total<br>suitable | Occupied<br>– no<br>dispersal | Occupied –<br>partial<br>dispersal |
| Portugal North | Current | 10,486,649 | 14 | 786 | 2,373,414 | 14 | 806 |
|  | 2021-2050 | 14,629,080 | 14 | 2,585 | 2,485,544 | 0 | 0 |
|  | 2051-2080 | 11,983,347 | 0 | 1,970 | 718,275 | 0 | 0 |
| Portugal South | Current | 2,379,798 | 51 | 2,830 | 1,727,140 | 51 | 2,792 |
|  | 2021-2050 | 108,753 | 43 | 6,084 | 0 | 0 | 0 |
|  | 2051-2080 | 18,008 | 41 | 12,625 | 0 | 0 | 0 |
| Spain | Current | 5,084,667 | 101 | 6,088 | 4,154,303 | 101 | 6,046 |
|  | 2021-2050 | 5,160 | 0 | 0 | 0 | 0 | 0 |
|  | 2051-2080 | 0 | 0 | 0 | 0 | 0 | 0 |

Under the most pessimistic scenario of climate change and ensemble model combination (A2 – mean), the existence of suitable climate was predicted to have disappeared completely by 2080 from two of the disjunct regions (southern Portugal & southern Spain), and have severely contracted in the third (northern Portugal) (Table 1). Considering dispersal ability, by 2050 it was predicted that dispersal limitations will have prevented the species from occupying any area of suitable climate across all regions.

### Invasive range and biogeographic projections

While ensemble models calibrated on the native range performed extremely well here according to model evaluation (sensitivity and specificity > 0.99 across all four EMs), these SDMs failed to predict the existence of suitable climate across the entire invasive range of this species in NW Europe (0 cells predicted as suitable across all model combinations). Within a biogeographical context, it was predicted that the number of areas with suitable climate for *R.p*. ssp. *baeticum* will rapidly decline within the present day Mediterranean biogeographic zone, while expanding within the Atlantic biogeographic region (Table 2). In some cases by 2050 (and in all cases by 2080), the majority of future climate suitable for *R.p*. ssp. *baeticum* was thus predicted to exist: A.) outside the current extant and documented historical range of this species; and B.) within geographical areas that fall outside the species’ present biogeographic region (Fig. 2).

**Figure 2.**
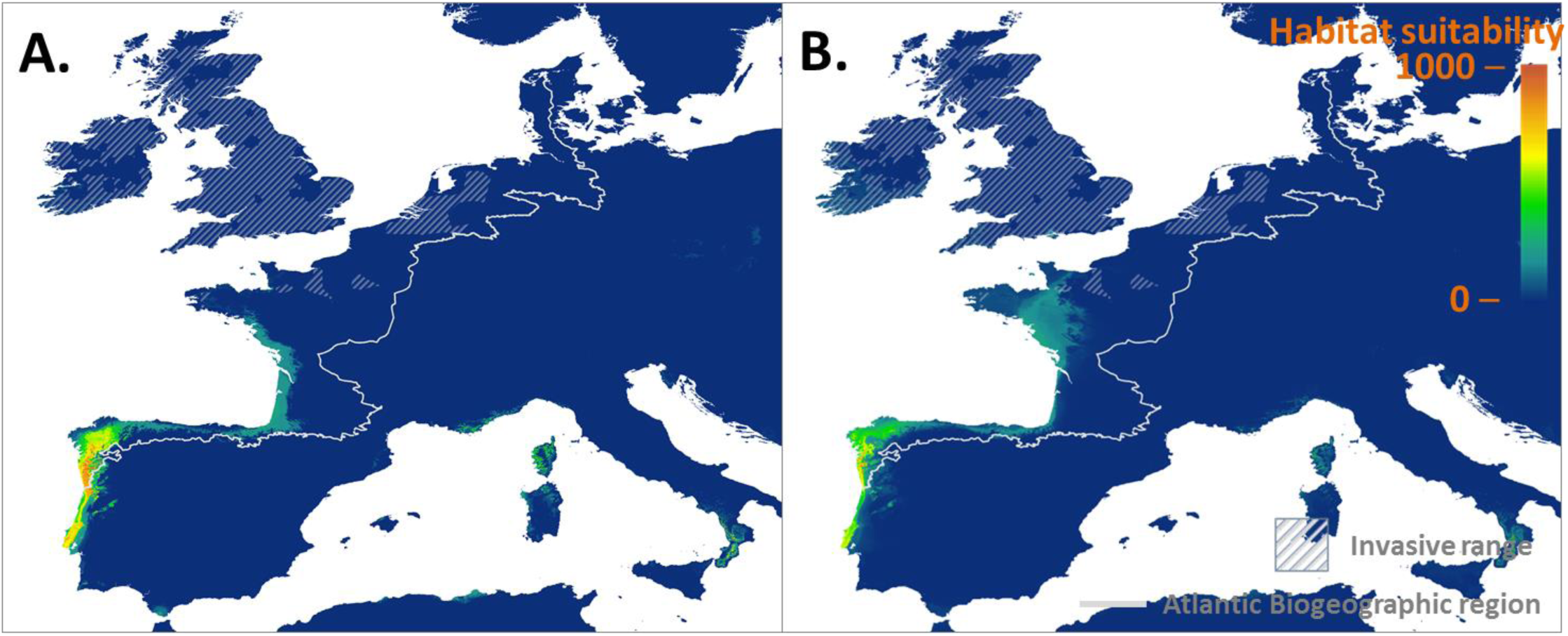
Areas of predicted climatic suitability for *R. ponticum* ssp. *baeticum* in the 2051–2080 time period, under A.) the most optimistic climate change scenario and ensemble model combination (B1 scenario – ca ensemble), and B.) the most pessimistic climate change scenario and ensemble model combination (A2 – mean ensemble). In both cases the majority of suitable climate is predicted to shift into areas of the present Atlantic biogeographic region.

**Table 2.** Number of 2 km^2^ cells possessing climatic suitability, and mean suitability score (from 0–1000) for *R. ponticum* ssp. *baeticum* within its extant native range, and per biogeographical zone on the Iberian Peninsula. Results are presented for the current climate period, and the most optimistic and pessimistic climate change scenario and ensemble model combination.

| Time period / Range | Extant native<br>range<sup>†</sup> | Mediterranean zone* | Atlantic zone* |
| --- | --- | --- | --- |
| No. 2 km<sup>2</sup> cells suitable / climatic suitability score (± SD) |  |  |  |
| **Current** |  |  |  |
| Current (EM ca) | 759 / 780 (± 264) | 6894 / 45 (± 154) | 742 / 69 (± 153) |
| Current (EM mean) | 486 / 603 (± 241) | 1591 / 42 (± 106) | 25 / 67 (± 97) |
| **B1 scenario/EM ca** |  |  |  |
| 2021-2050 | 293 / 405 (± 303) | 3756 / 32 (± 123) | 2368 / 158 (± 226) |
| 2051-2080 | 109 / 187 (± 235) | 2086 / 15 (± 83) | 3212 / 208 (± 246) |
| **A2 scenario/EM mean** |  |  |  |
| 2021-2050 | 0 / 163 (± 174) | 19 / 24 (± 63) | 121 / 145 (± 159) |
| 2051-2080 | 0 / 44 (± 36) | 1 / 18 (± 37) | 69 / 140 (± 138) |

<sup>†</sup> The present extant range was delimited by georeferencing distributional data from Perez Latorre and Cabezudo (2006) and Magos Brehm *et al.* (2008)
\*Confined to mainland Portugal and Spain only

### Identification of climate change microrefugia

The framework developed and applied as part of the current study successfully identified candidate sites potentially suited to translocation of *R.p*. ssp. *baeticum* for conservation purposes. A small number of 2 km cells in N Portugal and NW Spain (i.e. outside of the species’ historical distribution/present biogeographic zone) were identified as microrefugia most suited for potential translocation (Fig. 3). These cells indicated a relatively high level of accordance (83.3 %) in prediction of climatic suitability across the 24 different layers. As such, these sites were deemed to be temporally stable under a range of climate change scenarios, and robust to statistical uncertainties associated with the ensemble modelling techniques. However, as the species does not presently occupy these cells, and these do not possess climatic suitability under ‘current’ conditions, these sites represent future emergent microrefugia only (i.e. Fig. 1C).

**Figure 3.**
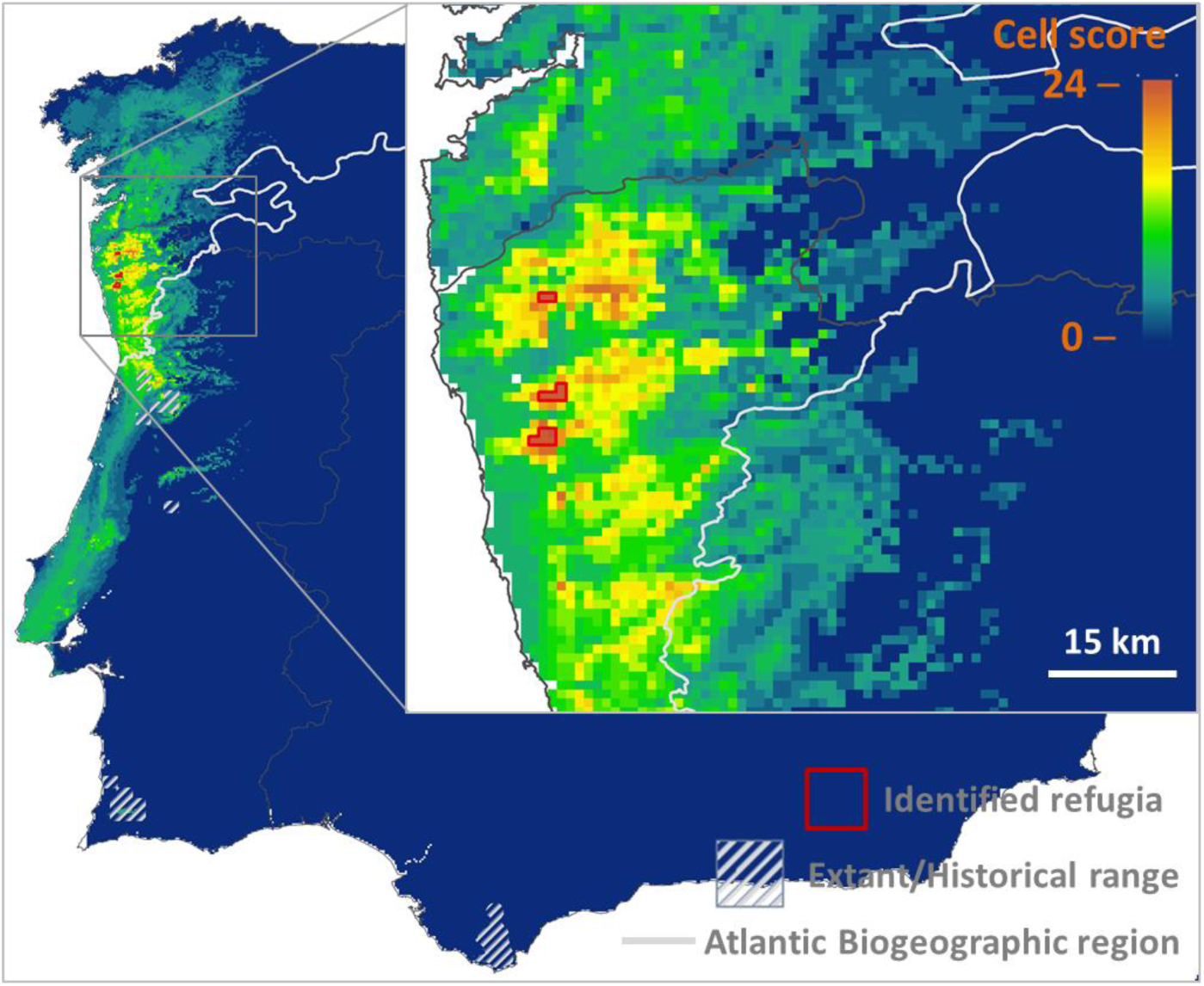
Identified putative climate change microrefugia for *R. ponticum* ssp. *baeticum* potentially suited to translocation. High scoring cells indicate areas with high accordance in predicted climatic suitability (between the various climate change scenarios and ensemble models employed). Efforts aimed at conserving this taxon from extinction by means of assisted colonization would hence most likely require translocation into microrefugia outside of the species’ extant (and known historical) range, and present

## DISCUSSION

### Migration and survival under climate change

Our results suggest that there is a strong likelihood of climate-induced extinction of *R. ponticum* ssp. *baeticum* in its native range under climate change. This risk is due to loss of suitable climate in areas of current occupation, coupled with an inability to migrate into future suited areas. Only under the most optimistic scenario (B1 climate change scenario – committee averaging EM) was it predicted that the species could occupy small areas of future suitable climate within the vicinity of its extant Portuguese range. In contrast, no occupation of future suitable climate was predicted anywhere in the most pessimistic scenario (A2 climate change scenario – mean EM). There was nonetheless consensus between scenarios in prediction of range collapse in this species’ disjunct Spanish range, where it is Red-listed nationally (Moreno, 2010). While our simulations of propagule dispersal for this species assumed full rates of germination and transition between life stages, in reality it is known that successful recruitment of new juveniles is extremely rare in the native range (Mejías et al., 2002). An almost categorical lack of natural recruitment would therefore best equate to our highly conservative predictions of migration ability based on ‘no dispersal’ (Table 1), and affirm the likelihood that climate change impacts will follow a path closer to our most pessimistic of trajectories.

Explanation as to why SDMs performed well according to models calibrated on the native range, yet did not predict suitable climate in the invaded range, may relate to the high specificity of *R.p. baeticum* to relict native habitat. Although climatic niche shifts are rare in the invasion processes (Petitpierre et al., 2012), they are known to occur. It is thus evident that *R.p. baeticum* possesses a larger niche breath than can be predicted from the limited range of environmental conditions occupied in the native range. Such cases of limited knowledge of a species’ ecology are highlighted as the biggest limit to implementation of translocation plans (Hancock & Gallagher, 2014). Nevertheless, while unrecognized niche breath or plasticity may lead to failure in predicating invasive capacity in novel geographic areas, such circumstances would also lead to more conservative estimates of sites suitable for translocation.

### The invasive alien/threatened native juxtaposition

Opinions currently differ as to the validity of translocation as a conservation strategy, as well as to the usefulness of a ‘native’ versus ‘alien’ duality in a future world where extinction due to migrational lag may be much more common-place (McLachlan et al., 2007; Warren, 2007; Ricciardi & Simberloff, 2009a; Ricciardi & Simberloff, 2009b; Warren, 2009; Minteer & Collins, 2010; Schwartz et al., 2012; Albrecht et al., 2013; Ahteensuu & Lehvävirta, 2014). Hence given their unique juxtaposition, climate-threatened invasive alien species (IAS) offer more than just hypothetical examples in this context. Given the uncertain survival of *R.p*. ssp. *baeticum* in its native range under climate change based on evidence presented in this study, this species could arguably qualify as a candidate for translocation in accordance with current guidelines (IUCN/SSC, 2013). However, as is the case for other threatened IAS (Suppl. Table 1), there exists a paradox in how best to approach practical conservation in these cases; including what role (if any) invasive populations may play at present and under future climate change. While eradication may for now continue as a goal within the introduced range, if risk of native extinction continues to intensify (as predicted here for *R.p. baeticum)*, the appropriateness of this strategy must at some point be called into question. Future actions could therefore include reintroduction of individuals or propagules into the native range, and/or acceptance of invasive populations as ‘climate-refugee’ species (Thomas, 2011). How realistic a prospect is a paradigm shift towards acceptance of such taxa as pre-translocated species, either now or in the future? As previously highlighted (Ohsawa & Jones, 2017), such perceptions may differ largely between stakeholders, rendering the general acceptance or application of such ideas challenging. How climate change will impact threatened IAS in their invasive range may ultimately steer such considerations, as future habitat suitability may be altered to levels which safely contain invasive spread, or even lead to invasive range contractions.

### Translocation under biogeographic constraints

While the theoretical aim of translocation is straight-forward, conceptual challenges are posed in complex cases such as for narrow or relict endemics like *R.p. baeticum*. Even in present times, such endemics may occupy regionally rare microclimates already under siege (Moreno, 2010). Further confounding this challenge is the fact that the present range of many endemics may also coincide with their known historic range – e.g. as due to long-term isolation in relict habitat, or a relatively recent evolutionary emergence (Lavergne et al., 2004; Hopper, 2009). Therefore translocations which are implemented outside of these areas are deemed as ‘introductions’, and this practice is not universally accepted (Ricciardi & Simberloff, 2009a; Ricciardi & Simberloff, 2009b; Bucharova, 2017). However, as evidenced by our results for *R.p. baeticum*, together with past findings on this species (Erfmeier & Bruelheide, 2010, 2011; Stout et al., 2015), translocation both outside this species’ present biogeographic zone, and extant/historic area of occurrence, represents the most feasible chance for its survival.

Our results indicated that although the application of a biogeographic constraint to the selection of translocation sites for *R.p. baeticum* would have evidently minimized invasive risk, strict interpretation of this principle would undoubtedly hamper efforts for conservation – especially when areas possessing long-term stability were readily identified in an adjacent biogeographic region a short distance from the extant range. Strict biogeographic constraints may similarly possess little relevance for many relict endemics, which often occupy microclimates that contrast sharply with the prevailing biogeographical zone. Furthermore, the usefulness of present day biogeographic delineations in guiding translocation may diminish the further in time projections are made, as such delineations are also set to shift rapidly under climate change (Williams et al., 2007; Mahlstein et al., 2013).

Hence based on the above considerations, we propose that where an unequivocal need exists, consideration should be afforded to implementing translocation outside of a species’ present biogeographical zone. Some authors have suggested to broaden the definition of a species’ ‘historic’ or ‘indigenous’ range to include any area once occupied in the past (Jørgensen, 2011). For *R. ponticum*, inter-glacial pollen records in fact indicate a past distribution of this species throughout much of the European continent (Cain, 1944; Jessen, 1948; Jessen et al., 1959; Martinetto, 2009), where it is presently considered non-native. Whether or not this former distribution should force reconsideration of this label is hence a subjective matter. While it is apparent that care must underpin any such interpretation of historic (or ‘historical’) range (Dalrymple & Moehrenschlager, 2013), successful identification of such areas could at least in theory guide and enhance the legitimacy of translocation outside of a species’ present biogeographical zone. That is provided a genuine need has been empirically established, and that such areas indeed possess current and future-stable climate.

### Identification of climate change microrefugia

We here developed and applied a framework for identification of climate change microrefugia robust to statistical uncertainty (based on consensus between ensemble models), and climatic uncertainty (between climate change scenarios). This framework is highly flexible to adaptation for additional inputs, which may depend on the level of robustness sought. For example, we here employed 24 different climate change scenario layers derived from one global circulation model (GCM) and two future time periods, but these inputs could be expanded to consider accordance between multiple other GCMs and finer temporal resolution, as desired. This framework is similar to that developed by Alagador et al. (2014), who worked on the setting up of protected areas for several species and on a limited budget. Yet, our framework focuses on detection of microrefugia at different time scales, and better matches the needs of single-species protection projects.

While this framework is relatively straightforward in principle, we believe it represents a robust tool to assist practical identification of climate change microrefugia and decision-making for purpose of conservation translocation. As translocations are generally considered the last line of action for averting species extinction, there is therefore significant pressure on practitioners to make informed and timely management decisions. However, the relative urgency and uncertainty at which such decision-making must frequently be undertaken means that due consideration has not always been afforded to the long-term suitability of recipient sites, especially where species decline is (at least in part) climate-induced. Our framework can therefore assist for such purposes, and given its temporal and spatially explicit nature, can help tackle the questions of ‘whether’, ‘where’ and ‘when’ translocation could or should take place. While identified candidate sites may not be numerous (i.e. in comparison to where a single modelling technique or climate change scenario is used), a crucial benefit is that identified sites possess high temporal stability and robustness to uncertainties, which may therefore minimize risk of failure. As differentiated from other frameworks for translocation (cf. Ferrarini et al., 2016), this strategy can be considered to take a ‘no regrets’ approach.

Application of this framework for *R.p. baeticum* revealed that this species does not presently occupy (or is likely to migrate into) long-term stable microrefugia. We were successful in identifying several small areas of putative future stable climate potentially suited for translocation by 2021–2050. While the temporal resolution of climate periods used in this study was coarse, shorter consecutive time intervals (e.g. 5–10 year periods) could afford greater accuracy as to forecasting when stable microrefugia may emerge and translocation ideally initiated. Such information is of critical importance for present contingency planning for climate-endangered species (Hunter, 2007), before climatic suitability in the extant range may undergo more serious decline. Where future emergent microrefugia are selected as recipient sites for translocation upon further scrutiny (e.g. of micro- and mesoscale climate and biotic factors), management action can hence be implemented in advance of planned establishment, and could include development of various habitat components to help ensure long-term survival.

## Acknowledgements

The authors wish to thank Juan Arroyo and Erin Jo Tiedeken for assistance locating native populations in the field, Mark Brown for helpful discussions in conceiving this study, Yvonne Buckley for providing input on an early draft, and Camila D’ Bastiani for research assistance. This work was funded by Science Foundation Ireland (10/RFP/EOB2842, to J.S.).

## Supplementary material

**Suppl. Table 1.**
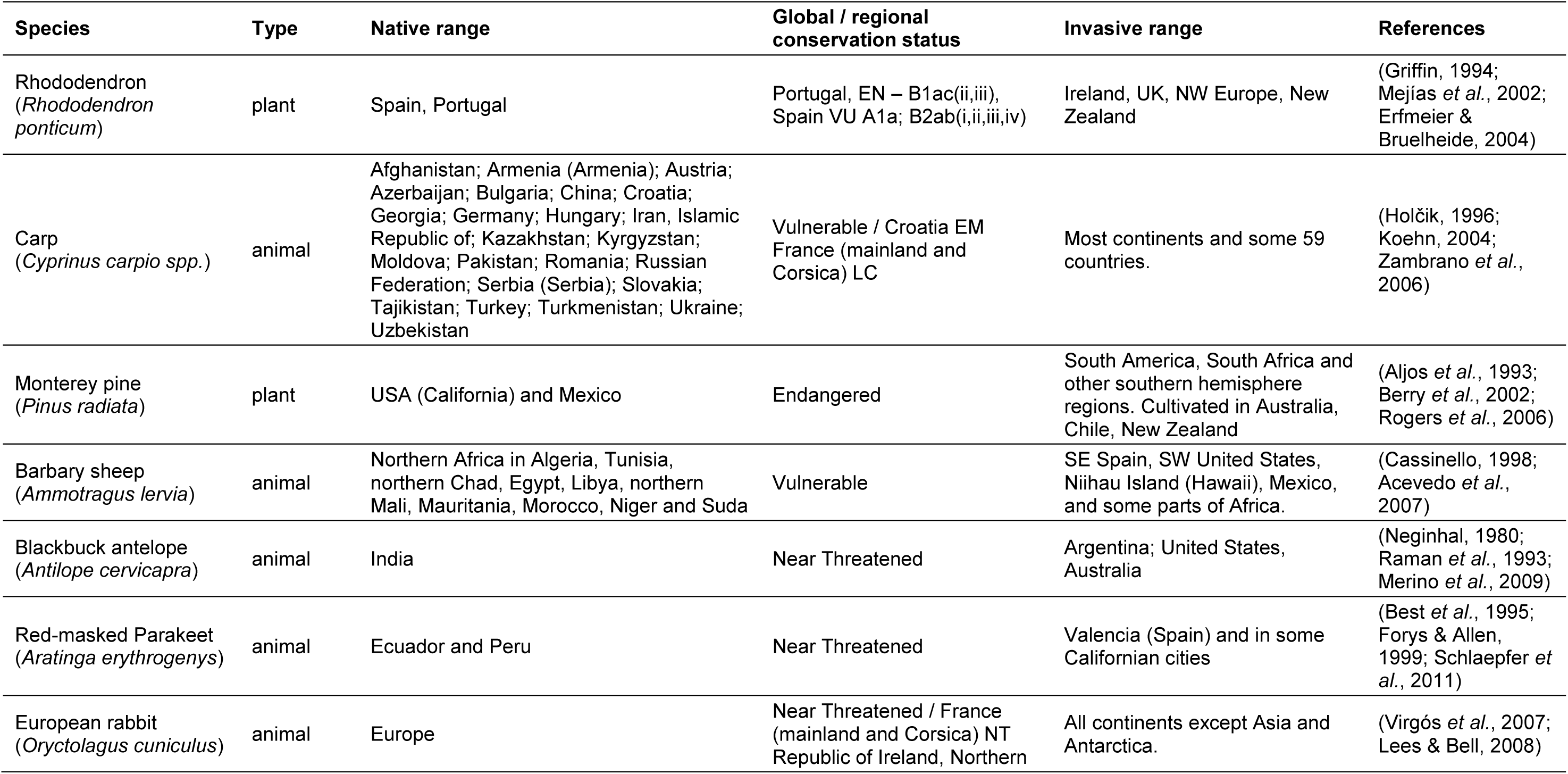
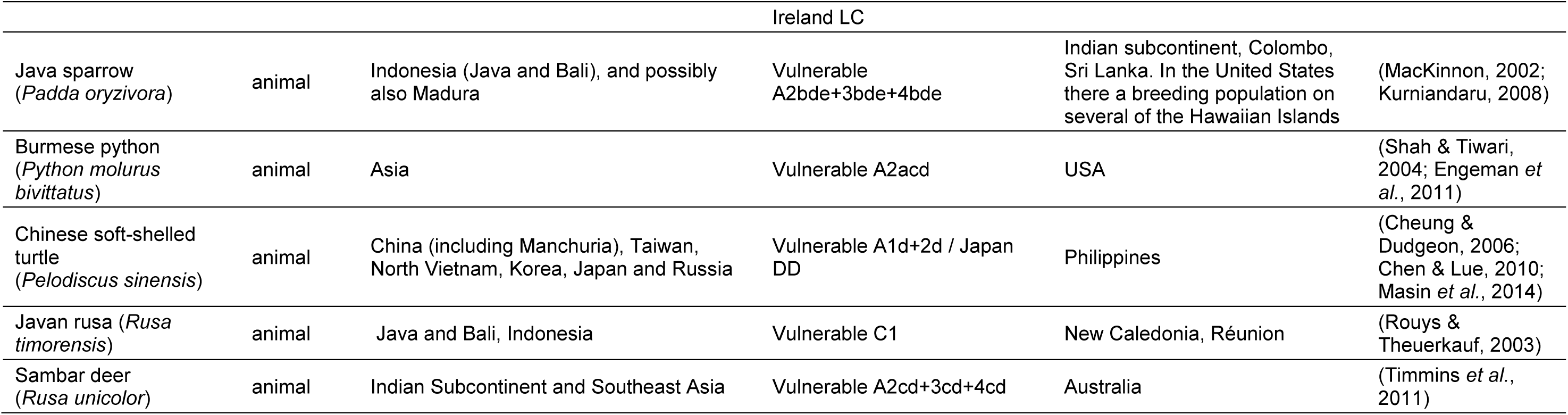
Invasive alien species (IAS) which are threatened in their native range. An initial list of 25–30 species loosely fitting these criteria were narrowed down to the 12 presented here (10 animal, 2 plant species) based on combination of their native conservation status and reports of their invasion intensity (which included a naturalised or colonized status across at least some parts of the introduced range). Representative literature sources, providing further details or anecdotal reports, are presented for each species.

**Suppl. Table 2.**
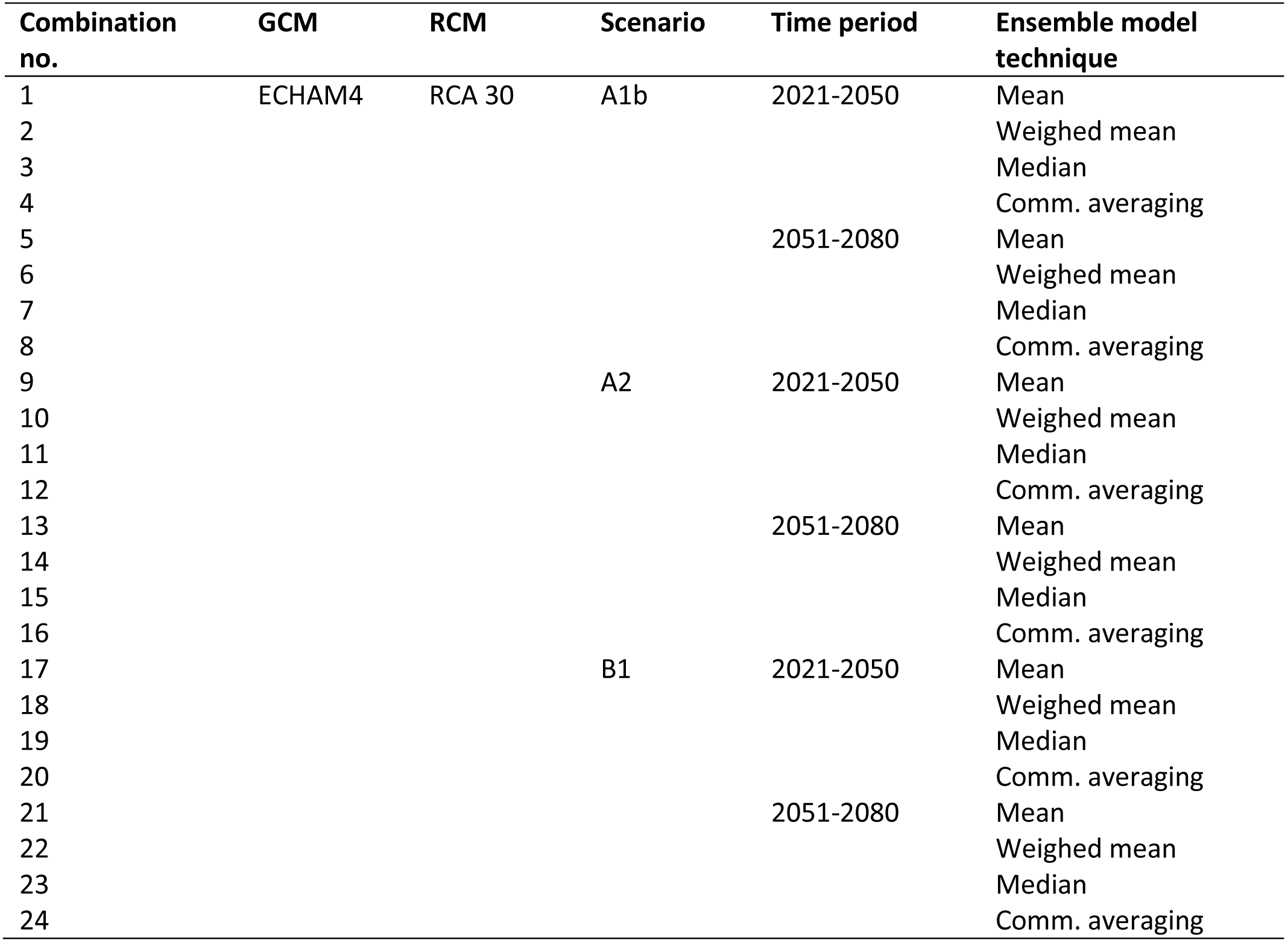
The 24 climate change layers utilised in this study, based on climate change scenario, time period, and the ensemble model technique employed.

| Combination no. | GCM | RCM | Scenario | Time period | Ensemble model technique |
| --- | --- | --- | --- | --- | --- |
| 1 | ECHAM4 | RCA 30 | A1b | 2021-2050 | Mean |
| 2 |  |  |  |  | Weighed mean |
| 3 |  |  |  |  | Median |
| 4 |  |  |  |  | Comm. averaging |
| 5 |  |  |  | 2051-2080 | Mean |
| 6 |  |  |  |  | Weighed mean |
| 7 |  |  |  |  | Median |
| 8 |  |  |  |  | Comm. averaging |
| 9 |  |  | A2 | 2021-2050 | Mean |
| 10 |  |  |  |  | Weighed mean |
| 11 |  |  |  |  | Median |
| 12 |  |  |  |  | Comm. averaging |
| 13 |  |  |  | 2051-2080 | Mean |
| 14 |  |  |  |  | Weighed mean |
| 15 |  |  |  |  | Median |
| 16 |  |  |  |  | Comm. averaging |
| 17 |  |  | B1 | 2021-2050 | Mean |
| 18 |  |  |  |  | Weighed mean |
| 19 |  |  |  |  | Median |
| 20 |  |  |  |  | Comm. averaging |
| 21 |  |  |  | 2051-2080 | Mean |
| 22 |  |  |  |  | Weighed mean |
| 23 |  |  |  |  | Median |
| 24 |  |  |  |  | Comm. averaging |

**Suppl. Table 3.** Climate variables and codes used. Variables in bold were used to build the models in the current study.

| **Code** | **Climate Variable** |
| --- | --- |
| BIO1 | Annual Mean Temperature |
| BIO2 | Mean Diurnal Range (Mean of monthly (max temp - min temp)) |
| BIO3 | Isothermality (BIO2/BIO7) (* 100) |
| BIO4 | Temperature Seasonality (standard deviation *100) |
| **BIO5** | **Max Temperature of Warmest Month** |
| BIO6 | Min Temperature of Coldest Month |
| **BIO7** | **Temperature Annual Range (BIO5-BIO6)** |
| BIO8 | Mean Temperature of Wettest Quarter |
| BIO9 | Mean Temperature of Driest Quarter |
| BIO10 | Mean Temperature of Warmest Quarter |
| **BIO11** | **Mean Temperature of Coldest Quarter** |
| BIO12 | Annual Precipitation |
| BIO13 | Precipitation of Wettest Month |
| BIO14 | Precipitation of Driest Month |
| BIO15 | Precipitation Seasonality (Coefficient of Variation) |
| **BIO16** | **Precipitation of Wettest Quarter** |
| **BIO17** | **Precipitation of Driest Quarter** |
| BIO18 | Precipitation of Warmest Quarter |
| BIO19 | Precipitation of Coldest Quarter |
| **ETP (PET)** | **Potential Evapotranspiration (annual mean)** |

**Suppl. Figure 1.**
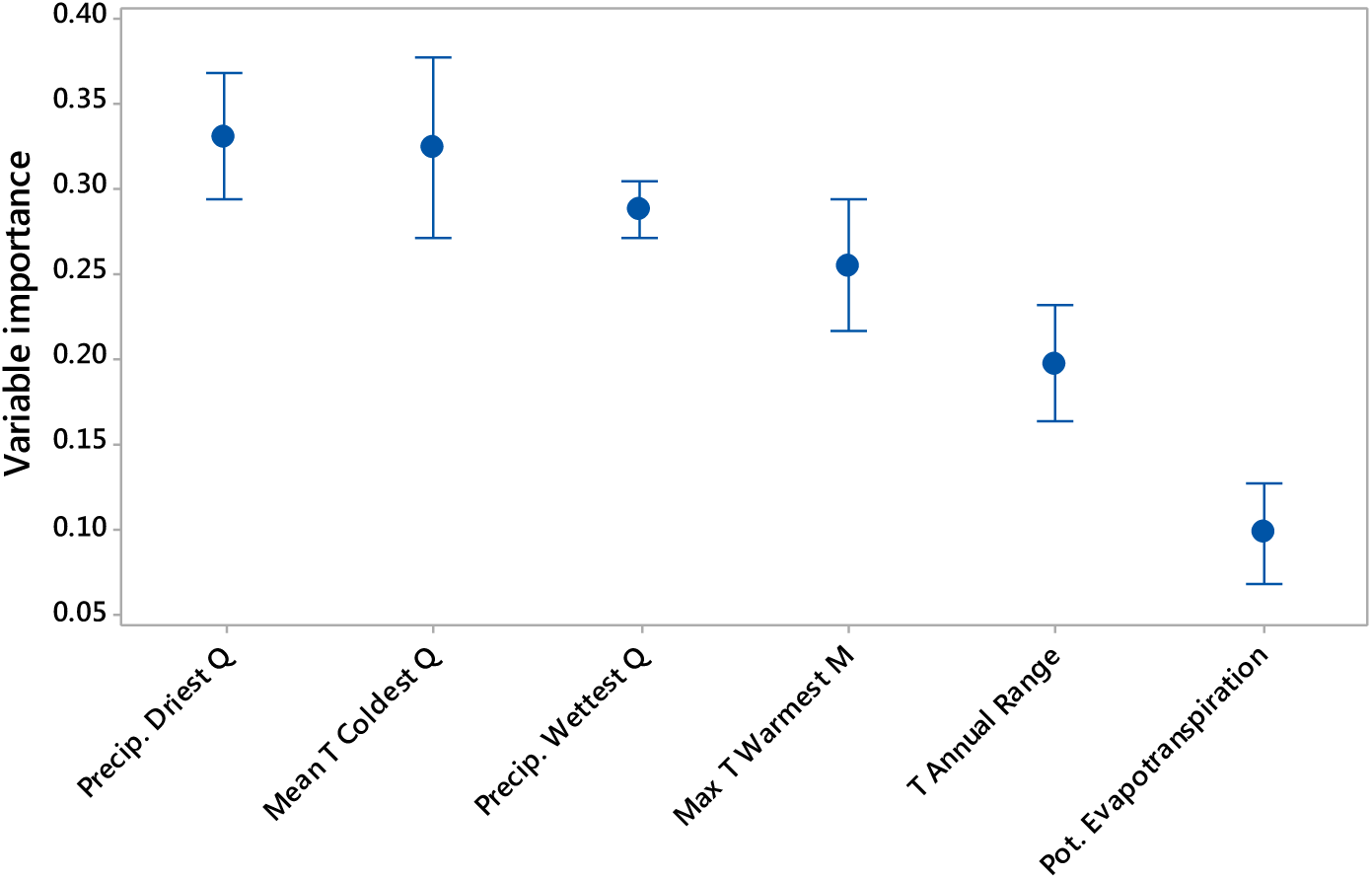
Variable importance (mean ± SE) for the six environmental variables used in modelling. These values represent the mean of 36 replicates, in which three different sets of pseudo-absences were used within three evaluation runs for each model type (GBM, GAM, RF, MAXENT).

